# RNA fragment assembly with experimental restraints

**DOI:** 10.1101/2021.02.08.430198

**Authors:** Grzegorz Chojnowski, Rafał Zaborowski, Marcin Magnus, Janusz M. Bujnicki

## Abstract

We present RNA Masonry, a computer program and a web service for a fully automated assembly of RNA fragments into geometrically plausible models fulfilling user-provided secondary structure constraints and restraints on tertiary contacts and Small Angle X-ray Scattering (SAXS) data. We illustrate the method description with its recent application to structural studies of viral RNAs with SAXS restraints. The program web server is available at http://iimcb.genesilico.pl/rnamasonry.

## 1. Introduction

Non-coding RNAs (ncRNAs) are involved in the regulation of many cellular processes. New families of ncRNAs are being continuously discovered (Kalvari et al. 2018; Boccaletto et al. 2018). In principle, the function of newly characterized ncRNAs could be deciphered from their structure in an approach that proved very successful for proteins (Miao et al. 2015; Doudna 2000). The experimental structure determination of ncRNAs (e.g., by X-ray crystallography, nuclear magnetic resonance, or cryo-electron microscopy) is however difficult, and only a very small fraction of ncRNA families have high-resolution structures available for at least one member (99 out of 3,016 according to Rfam 14.1). Computational structure prediction methods offer an alternative to experimental structure determination, but the purely theoretical models are often of limited accuracy. One promising approach is the computational modelling with the use of restraints derived from low-resolution experimental data (Magnus et al. 2014). To address this issue we developed RNA Masonry, a computer program for fully automated modelling of RNA molecules by assembly of recurrent 3D motifs, with the use of low-resolution restraints. The 3D motifs (further referred to as fragments) are retrieved from the RNA Bricks database (Chojnowski, Walen, and Bujnicki 2014), which catalogues recurrent substructures observed in experimentally-determined, high-resolution RNA structural models available in the Protein Data Bank (Berman et al. 2000).

RNA Masonry exploits hierarchical organization of RNA molecules, which are composed of 3D motifs defined at a secondary structure level; double-stranded helices, single-stranded fragments, and various types of loops. During the model assembly, the motifs are used as a whole, which strictly preserves the input secondary structure. The program accepts restraints for long-range tertiary interactions. Additionally, the model building can be restrained with a goodness-of-fit to the experimental small angle X-ray scattering (SAXS) data, which is calculated using FoXS (Schneidman-Duhovny et al. 2013) or CRYSOL (Svergun, Barberato, and Koch 1995).

There are other methods available that can assemble RNA 3D structures from fragments of experimentally determined structures (Jossinet, Ludwig, and Westhof 2010; Popenda et al. 2012) one of them designed specifically for the purpose of modelling with SAXS restraints (Gajda et al. 2013). To the best of our knowledge, however, RNA Masonry is the only tool that combines statistical potential with experimental restraints and uses a regularly updated database of fragments (RNA Bricks is updated weekly, http://genesilico.pl/rnabricks2).

## 2. Materials and methods

The program provides two basic functionalities: 1) *de novo* fragment assembly and 2) RNA model refinement. The input is either RNA sequence with the secondary structure or a preliminary RNA 3D structural model in the PDB or mmCIF format. The program can use SAXS data as an additional restraint in any format recognized by FoXS and CRYSOL.

### 2.1 Input processing

If the input is an RNA atomic model, the program annotates its secondary structure with ClaRNA (Waleń et al. 2014) which is further processed analogously to a secondary structure given directly on input for *de novo* modelling.

Pseudo-knots are removed from the input secondary structure using K2N (Smit et al. 2008) and stored as additional restraints. The pseudo-knot free secondary structure is subsequently decomposed into structural motifs and encoded as an RNA motif-graph introduced in the RNA Bricks database. Briefly, the graph nodes correspond to RNA structural motifs (double-stranded helices, loops, and single-stranded fragments). The graph edges denote nucleotides shared by two neighbouring motifs. Owing to the absence of pseudo-knots in the basic data structure the motif graphs are trees, albeit not necessarily rooted. Pseudoknots are nonetheless enforced at the level of structural restraints.

### 2.2 Selecting RNA 3D fragments

After building the motif-graph, which is a basic data structure used in RNA Masonry, 3D fragments from RNA Bricks database are assigned to each of its nodes following the secondary structure constraints. The fragments are selected based exclusively on the canonical secondary structure. No sequence information is taken into account. If the number of fragments assigned to a node is fewer than a user-defined threshold (10 by default) additional fragments are generated by circular permutations (e.g., by changing ends of symmetrical, internal loops). If needed, new fragments are also automatically built from smaller ones by introducing insertions using ModeRNA (Rother et al. 2011). Finally, each fragment is mutated to the target sequence. Any steric conflicts, unsatisfied pseudo-knot restraints, or implausible geometries introduced at this stage are ignored, but penalized later during fragment assembly steps by the SimRNA (Boniecki et al. 2016) scoring function.

### 2.3 Starting RNA model assembly

If RNA 3D structure is given as an input, it is used as a starting model without any further modifications. Otherwise, the program assembles a starting model with a possibly small number of severe clashes, which readily increases the convergence of the subsequent refinement step.

In principle, the aim is to find a configuration of fragments, assigned to each of the motif-graph nodes, which minimizes the number of clashes in a complete model. A clash occurs when two glycosidic nitrogens (N1 for pyrimidines, N9 for purines) are closer than 6 Å apart. To reduce the combinatorial complexity of the task, the assembly process is initiated from the terminal elements of the structure (the motif graph “leaves”) and proceeds iteratively by adding a single node at a time. At each step, from all possible partial models only statistically relevant representatives are selected as described by Hajdin et al. (Hajdin et al. 2010).

### 2.4 Model optimization

The starting model is optimized in a replica exchange Monte-Carlo simulation with a geometric distribution of temperature levels. We use a scoring function that combines SimRNA statistical potential (Boniecki et al. 2016) and SAXS curve goodness-of-fit (if given on input). The long-range tertiary contact restraints can be also included (e.g., to indicate pseudoknots). In a single step, a random motif-graph node is selected. Next, from a set of fragments assigned to the node a random one is selected and inserted into corresponding position in the model. For all motifs except terminal loops, this operation changes conformation of a whole domain in the RNA model. The user-provided secondary structure is preserved at each step. The lowest-score model constructed during the entire simulation is returned to the user.

## 3. Structural studies of viral RNAs with SAXS restraints

We used an early version RNA Masonry for modelling adenovirus virus-associated RNA (VA-I) structure with SAXS restraints (Dzananovic et al. 2017). Interestingly, a crystal structure of VA-I has been recently determined and turned out to be inconsistent with the results of scattering experiments with goodness-of-fit parameter (X^2^) estimated using CRYSOL of 2.8 (Hood et al. 2019) (Fig. 1A). We used RNA Masonry to optimize the fit of the structure to the SAXS curve, while preserving its secondary structure. The resulting model fits better not only to the experimental data (X^2^=1.5), but also to the *ab initio* reconstruction obtained independently using DAMMIF (Franke and Svergun 2009) (Fig. 1B). This result supports a hypothesis that in solution and in the absence of crystal-lattice constraints VA-I may be flexible and sample additional conformations, possibly at the expense of the pseudoknot disruption (Hood et al. 2019).

**Figure 1.**
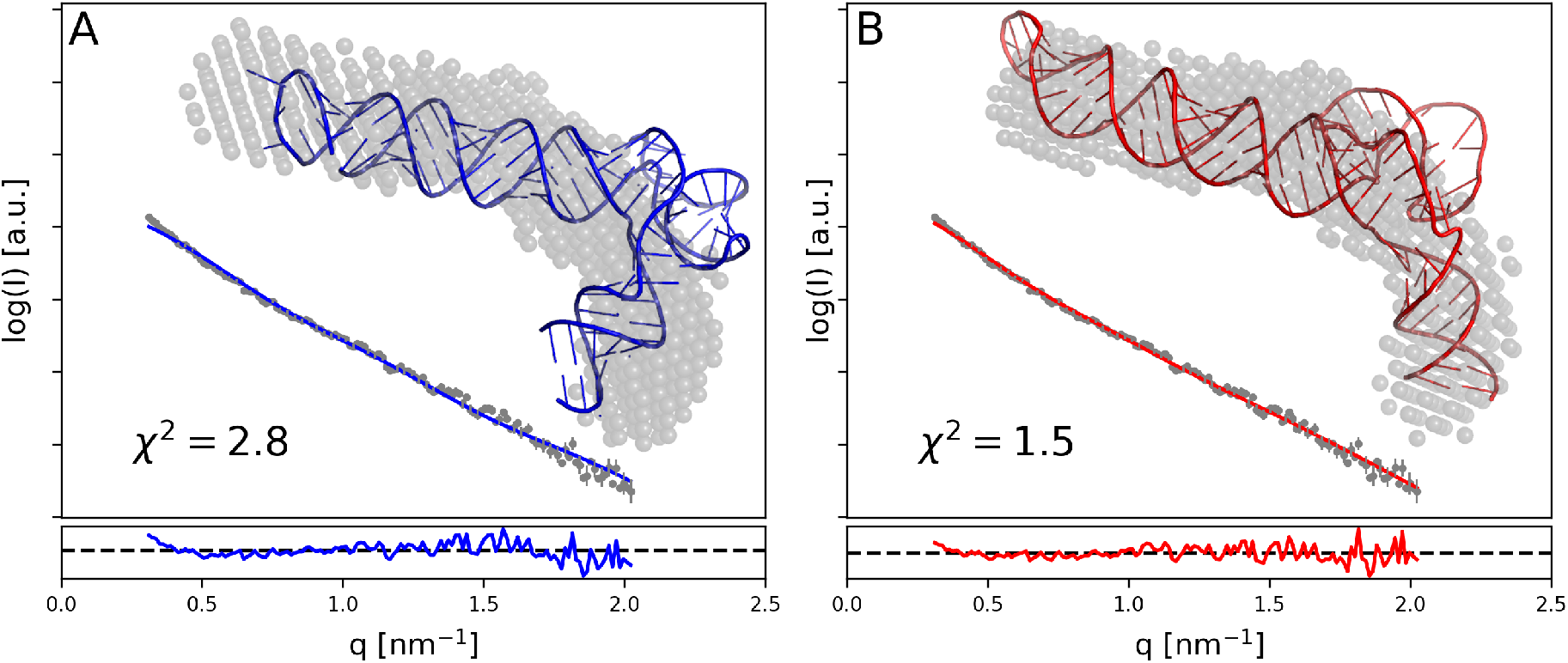
Crystal structure of adenovirus virus-associated RNA (A) was observed to have a conformation inconsistent with results of earlier SAXS experiments. RNA Masonry was used to optimize the fit to experimental SAXS data preserving the crystal structure secondary contacts (B). An ab-initio model obtained with DAMMIF is shown with grey spheres. Residuals of model-calculated and experimental SAXS curve fit are shown in the bottom panels.

## 4. Implementation

RNA Masonry and all utility programs were implemented in Python 2.7 and C++ with an extensive use of routines from the Computational Crystallography Toolbox (cctbx) (Grosse-Kunstleve et al. 2002), Numpy (Oliphant 2006), Scipy (Virtanen et al. 2020), NetworkX (Hagberg, Swart, and S Chult 2008), Biopython (Cock et al. 2009), ClaRNA, ModeRNA, and K2N libraries. Migration to Python version 3, which requires an update of some of the RNA Masonry dependencies, is planned in due course. The web server engine used in this work is a part of rna-tools toolbox (Magnus et al. 2020) and can be freely reused for new applications.

## 5. Conclusions

RNA Masonry is a computer program for fully automated modelling of RNA 3D structures by fragment assembly. The RNA structure representation used here significantly reduces the number of degrees of freedom, which in principle equals the number of joints (internal loops) in the target structure. The small number of degrees of freedom significantly reduces the modelling time (e.g., compared to SimRNA simulations). In the case of modelling with SAXS data it also reduces the risk of overfitting, given the limited number of unique observations that are generally available from SAXS experiments (Konarev and Svergun 2015). It must be stressed, that at the same time the presented approach is instrinctly unable to model fine details of RNA structures (e.g., non-canonical interactions) that are not encoded within the 3D motifs used for assembly. These, however, can be modeled using complementary high-resolution approaches (e.g., using QRNAS (Stasiewicz et al. 2019), SimRNA, or ROSETTA/FARFAR (Yesselman and Das 2016)).

The current RNA Masonry version selects RNA fragments for model-building based exclusively on canonical secondary structure constraints. It was observed, however, that the presence of certain motif types (e.g., kink-turn (Klein et al. 2001)) and coaxial stacking between adjacent helices can be reliably predicted from sequence (Cruz and Westhof 2011; Tyagi and Mathews 2007). Both these *a priori* information could be in principle used to restrain selection of fragments during model building in RNA Masonry to increase reliability of resulting models. We plan such an extension in further releases of the software.

## Acknowledgements

We thank Dr. Michal Boniecki (IIMCB) for sharing the SimRNA scoring function and Dr. Tomasz Waleń (University of Warsaw) for sharing ClaRNA source code.

